# Gene expression profiles of chemosensory genes of termite soldier and worker antennae

**DOI:** 10.1101/2022.07.25.501343

**Authors:** Ryohei H. Suzuki, Takumi Hanada, Yoshinobu Hayashi, Shuji Shigenobu, Kiyoto Maekawa, Masaru K. Hojo

## Abstract

Termite caste differentiation and social behavior are appropriately regulated by the chemical signals among individuals in each colony. Signal transduction is well known to be triggered by the reception of odorant molecules by some binding proteins in the antennae, after which, a signal is transmitted to chemosensory receptors. However, there is insufficient information on the role of chemosensory genes involved in signal transduction in termites. Here, we identified the genes involved in chemosensory reception in the rhinotermitid termite *Reticulitermes speratus*, and performed a genome-wide comparative transcriptome analysis of worker and soldier antennae. First, we identified 31 odorant-binding proteins (OBPs), and three chemosensory protein A (CheA) from the available genome sequence data. Thereafter, we performed RNA sequencing to compare the expression levels of OBPs, CheAs, and previously identified chemosensory receptor genes between workers and soldiers antennae. Of note, there were no receptor genes with significant differences in expression between castes. However, the expression levels of three non-receptor genes (*OBP, CheA*, and *Sensory neuron membrane protein*) were significantly different between castes. Quantiative polymerase chain reaction (qPCR) analysis using antennae and other head parts confirmed that these genes were highly expressed in soldier antennae. Finally, independent qPCR analysis showed that the expression patterns of these genes were altered in soldiers from different social contexts. The present results suggest that some non-receptor protein genes are involved in the social behaviors of termites.

## INTRODUCTION

Termites, one of the major type of social insects, have distinctive morphological and behavioral phenotypes (castes), including reproductives, soldiers, and workers (Wilson, 1971). Cooperation and division of labor are precisely achieved among colony members to maintain an elaborate social system. Determining the reason for the acquisition of a sophisticated social life is one of the main research goals in the field of evolutionary biology. Regulatory mechanisms of caste differentiation caused by interactions among colony members have focused on termites (Watanabe et al., 2014). Volatile candidate primer pheromones have been identified in some species (Matsuura et al., 2010; Mitaka et al., 2016; Tarver et al., 2009). However, it remains unclear how external environmental factors, including primer pheromones, influence internal physiological (e.g., hormone titer) changes during caste differentiation.

Generally, there are three receptor families involved in chemoreception in insects: Odorant receptors (ORs), Ionotropic receptors (IRs), and Gustatory receptors (GRs) (Nei et al., 2008; Croset et al., 2010). External factors, including pheromones, are usually recognized by these receptors (Clyne et al., 1999, 2000; Benton et al., 2019). Most genes encoding receptor proteins are expressed in sensory fibers in each sensillum (Koontz and Schneider, 1987; Gnatzy et al., 1984). The internal regions are filled with sensillar lymph, and hydrophobic chemicals bind to protein transporters and move to the appropriate receptors (Pelosi et al., 2016). ORs are the most commonly studied family of chemoreceptors. OR proteins are normally localized on the dendrites of odorant receptor neurons (OSNs) and they comprise seven transmembrane domains (Clyne et al., 1999, Vosshall et al., 1999). It is well known that the olfactory receptor co-receptor, Orco, is required for the function of each OR, and the OR/Orco complex is formed in chemosensory neurons (Larsson et al., 2004). The deletion of Orco disrupts the function of all ORs (DeGennaro et al., 2013; Larsson et al., 2004). Moreover, sensory neuron membrane proteins (SNMPs) were originally identified as abundant proteins on OSN dendrite membranes in moths (Rogers et al., 1997). SNMPs belong to the CD36 protein family and have been identified in several insect species (Vogt et al., 2009; Zhang et al., 2020; Zhao et al., 2020). One of the two subfamilies of SNMPs (Vogt et al., 2009) has shown to be expressed in antennal OSNs (Forstner et al., 2008; Gu et al., 2013; Blankenburg et al., 2019). These expression patterns may be involved in the functions of OR co-receptors and/or the transfer of ligands (including pheromones) to ORs (Gomez-Diaz et al., 2016).

In ants, more than 300 OR genes have been identified, and the OR subfamily with nine exons (known as 9-exons) has diverged among species (Zhou et al., 2012). The number of chemoreceptor families is much larger in ants than in other related hymenopteran lineages, suggesting that ants acquired complex chemical communication using various chemical substances detected by ORs (Gadau et al., 2012). Notably, in contrast to ants, IRs specifically diverged in the dampwood termite, *Zootermopsis nevadensis* (Terrapon et al., 2014). More than 400 IRs have been identified in the German cockroach, *Blattella germanica*, and IRs may be involved in specific chemical communication in both termites and cockroaches (Harrison et al., 2018).

In addition to receptors, many other protein families are involved in the transport of various chemical substances. The two major insect transporters are odorant-binding proteins (OBPs) and chemosensory proteins (CSPs), which are hydrophilic proteins with spherical shapes that have pockets containing hydrophobic substances (Pelosi et al., 2006). These pockets contain an alpha-helical structure and various numbers of disulfide bonds with conserved cysteine residues (Pelosi et al., 1995). CSPs have two disulfide bonds (Angeli et al., 1999), while most OBPs have three bonds (Leal, Nikonova & Peng, 1999; Scaloni et al., 1999), and some OBPs have an increased or decreased number of cysteines (known as C-plus and C-minus OBPs, respectively) (Xu, Zwiebel & Smith, 2003; Zhou et al., 2004; Lagarde et al., 2011; Spinelli et al., 2012). Most OBPs and CSPs are thought to be involved in the reception of chemical substances, including pheromones (Gracia et al., 2009). For example, in the carpenter ant, *Camponotus japonicus*, CSPs are bound to cuticular hydrocarbons (CHCs) and are involved in the recognition of colony members (Ozaki et al., 2005; Hojo et al., 2014). Additionally, other chemosensory proteins (known as Ches) have been identified in the chemoreception organs of *Drosophila melanogaster*, and have been classified into two subfamilies, CheA and CheB (Xu et al., 2002). Although the function of CheA is unclear, CheB may be involved in the reception of sex pheromones in flies (Xu et al., 2002; Touhara & Vosshall, 2009). Ches have also been identified in non-dipteran insects and are likely to be fast-evolving genes (Torres-Oliva et al., 2016).

In this study, we focused on the chemosensory genes of the termite, *Reticulitermes speratus*. The genome of this species has been sequenced and many chemosensory genes (31 ORs, 25 GR s, 92 IRs, 5 SNMPs, and 10 CSPs) have been identified (Shigenobu et al., 2022). Most chemosensory genes are probably expressed in the chemoreception organs, and the antenna is one of the most important reception organs. Indeed, antennal transcriptome analyses have been performed in the leaf-cutting ant, *Atta vollenweideri*, and the American cockroach, *Periplaneta americana*, and some receptor genes with differentially expression patterns between castes or sexes have been identified (Koch et al., 2013; Chen et al ., 2016). Consequently, we focused on the transcriptomes of the worker and soldier antennae in *R. speratus*. First, we identified binding protein genes (OBPs and CheAs) from *R. speratus* genome data (Rspe OGS1.0; Shigenobu et al., 2022). Second, we performed RNA-sequencing (RNA-seq) analysis using worker and soldier (female and male) antennal tissues, and compared the expression levels of chemosensory genes between castes and sexes. We verified the RNA-seq results using real-time quantitative polymerase chain reaction (qPCR) using gene-sepecic primers (Supplementary Table S1), and identified very small numbers of chemosensory genes with differentially expression patterns between castes. Finally, we performed the independent qPCR analysis to determine whether expression patterns of chemosensory genes were altered by different social contexts. Based on the data we obtained, we discuss the potential importance of certain transporter genes in the social behaviors of *R. speratus* workers and soldiers.

## RESULTS AND DISCUSSION

### Identification of *OBPs* and *CheA*

Using the *R. speratus* gene model (Rspe_OGS1.0), we identified 31 OBPs (*RsOBP1-31*). The number of OBPs was similar to the number of OBPs in *Z. nevadensis* (29 OBPs; Terappon et al. 2014; Table 1). In total, 19 RsOBPs were completely modeled, including full-length gene sequences with start and stop codons. Multiple alignments of these 19 RsOBPs with those of other insect species indicated that 16 were classic OBPs with six conserved cysteine residues, and the remaining three were Minus-C OBPs with conserved cysteine residues, according to the classification system used for insect OBPs (Hekmat-Scafe et al., 2002; Fan et al., 2011). Based on the RNA-seq *de novo* assembly data, nine “RsOBPs” have previously been reported (Mitaka et al., 2016). However, only one gene (“*RsOBP7*” reported by Mitaka et al., 2016) was identified in this study (*RsOBP22*), and the remaining “RsOBPs” could not be obtained from the genome sequence data. We identified the OBPs of *M. natalensis* from genome sequence data (21 genes) and performed molecular phylogenetic analysis. The results indicated that 30 of the 31 RsOBPs identified here showed sister-group relationships with those of termites [*M. natalensis* and/0r *Z. nevadensis* (total of 27 genes)] or cockroaches [*B. germanica* (total of three genes)] (Supplementary Fig. 1).

Thereafter, based on the *R. speratus* gene model (Rspe_OGS1.0), we identified three CheA genes (*RsCheA1-3*; Table 1). The number of CheAs identified in termites and cockroaches was different for each species, and the number of CheAs identified in *B. germanica* (more than 10) was much more than the number identified in termites (three to five genes; Supplementary Fig. 2). A previous study identified *CheB*-homologous genes from different orders of insects using *D. melanogaster CheB* sequences as a query (Torres-Oliva et al., 2016). However, *CheB*-homologous sequences were not obtained from the genomic or transcriptomic data of the termites and cockroaches analyzed here (*R. speratus, M. natalensis, Z. nevadensis*, and *B. germanica*). Molecular phylogeny analysis showed that all *CheAs* identified in termites and cockroaches were closely related to the *D. melanogaster CheA7a* gene sequence (Supplementary Fig. 2).

Overall, extant copies of both *OBPs* and *CheAs* in termites and cockroaches were orthologous among each species (Supplementary Figs. 1 and 2). These results suggest that gene duplication events may have occurred before the appearance of termites that diverged from a cockroach ancestor. Further analysis should be performed to determine whether there are differences in the function of each gene copy, especially between termites and cockroaches.

### RNA-seq analysis of worker and soldier antennae

We identified 540 differentially expressed genes (DEGs) between soldier and worker antennae (Fig. 1A, Supplementary Table S2). Of these genes, 234 and 306 were upregulated in workers and soldiers, respectively (Fig. 1B). In contrast, there were only one and five DEGs between male and female soldiers and workers, respectively (Fig. 1B). RNA-seq data showed that very small numbers (only four) of chemosensory genes identified from the genome were significantly highly expressed in soldier or worker antennae (Fig. 2, Supplementary Table S3–S9). High expression levels of three non-receptor genes [*RsOBP27* (gene ID: *RS009616*), *RsSNMP2* (*RS007977*), and *RsCheA1 (RS004717*)], were observed in soldier antennae, compared with worker antennae. One receptor gene [*RsIR75o* (*RS012794*)] was highly expressed in worker antennae. However, RsIR75o was not completely modeled and did not contain both the start and stop codons. Previous RNA-seq data indicated that *RsIR75o* (*RS012794*) was not expressed in any of the castes examined (reads per kilobase of transcript, per million mapped reads = 0; Shigenobu et al., 2022).

**Fig. 1.**
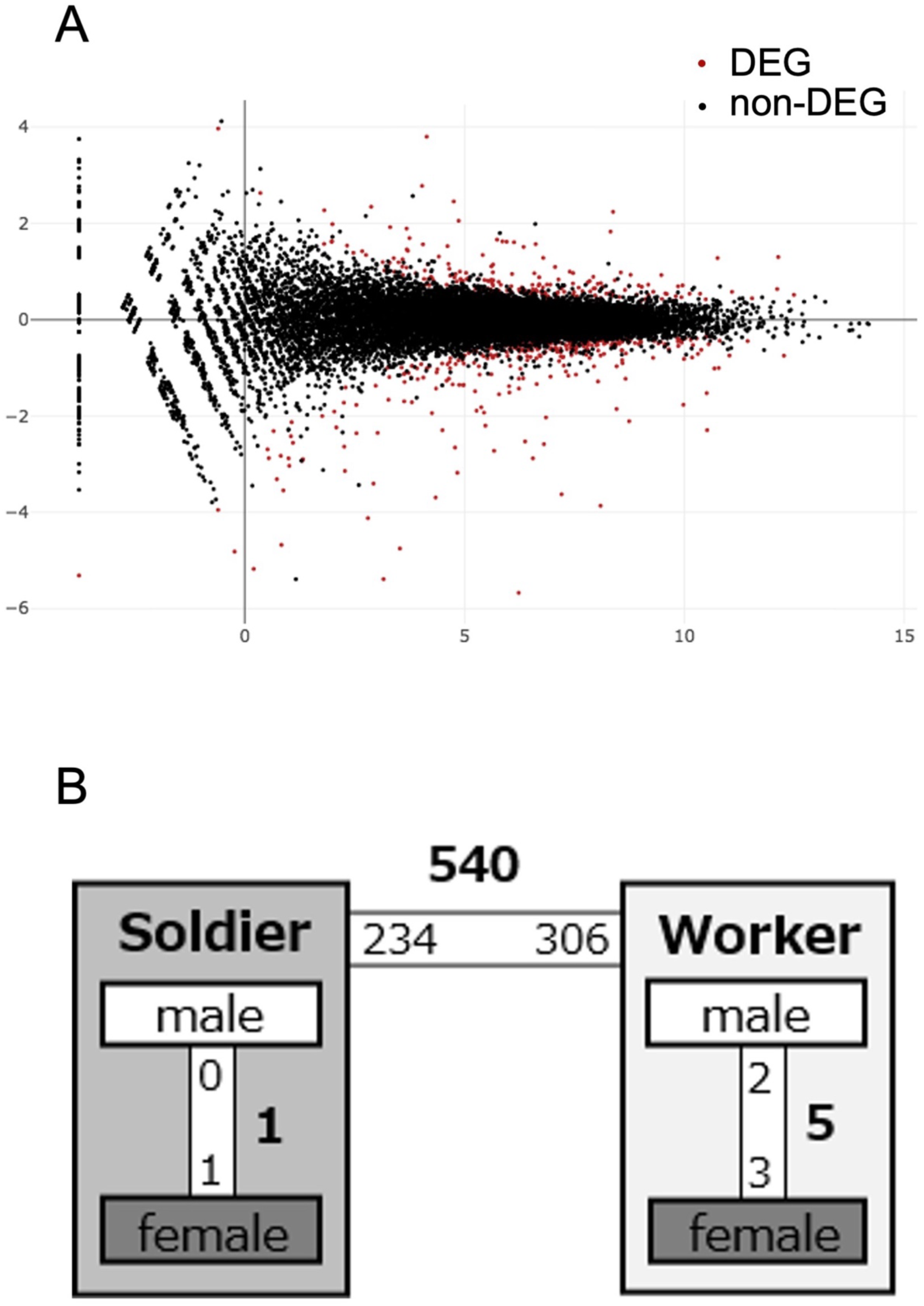
**(A)** MA plot of RNA-seq data obtained in this study. The x-axis indicates a log mean value of counts per millions (CPM). The y-axis indicates a log fold value. The color plots show significantly (red) and non-significantly (black) differentially expressed genes (DEGs; false discovery rate < 0.05). (**B)** The numbers of DEGs (bold) between each category. Small number indicate the numbers of upregulated genes in each category. The numbers of DEGs between soldiers and workers (540) was much larger than the number of DEGs between males and females (total six).

**Fig. 2.**
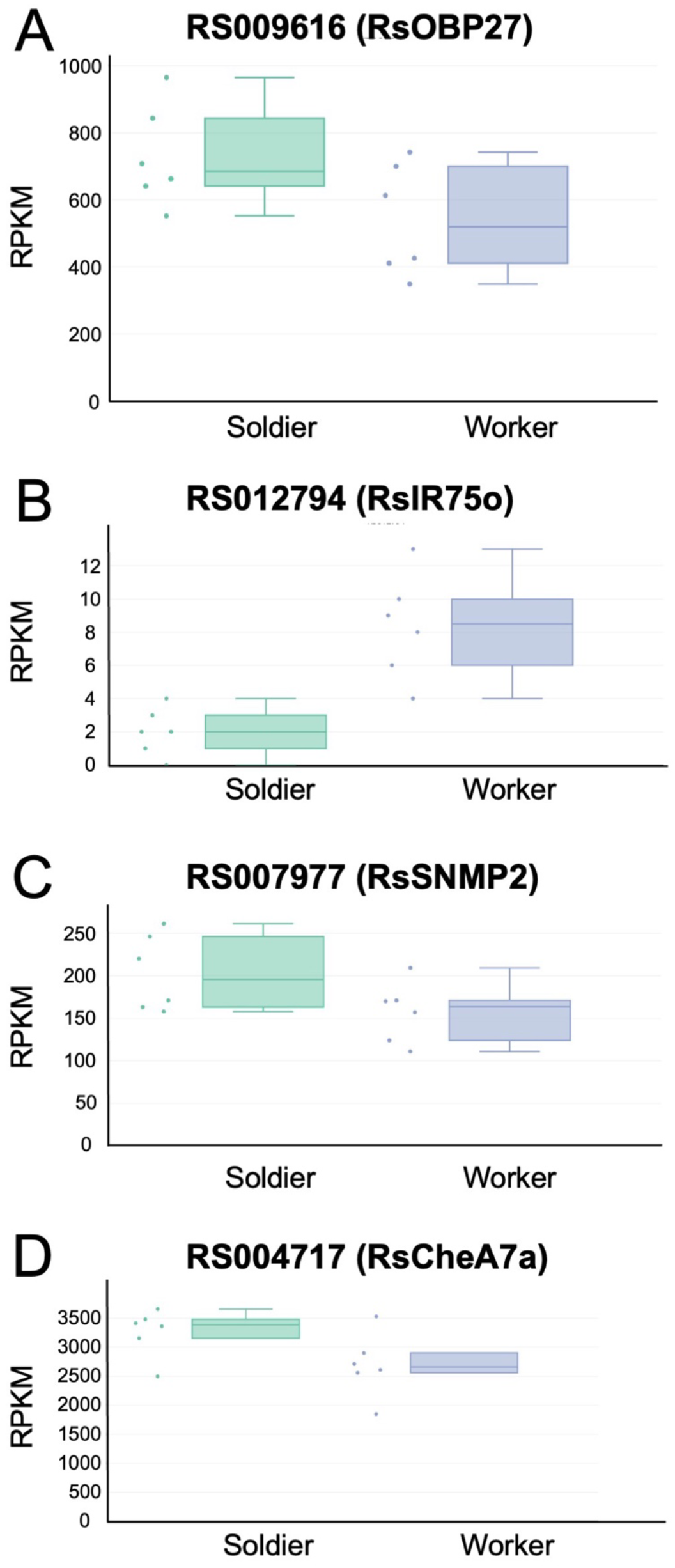
**(A)**-**(D)** Expression levels of **(A)** *RS009616* (*RsOBP27*), **(B)** *RS012794* (*RsIR75o*), **(C)** *RS007977* (*RsSNMP2*) and **(D)** *RS004717* (*RsCheA1*) between *Reticulitermes speratus* soldier and worker antennae. Expression levels are indicated as read per kilobase of transcript, per million mapped reads (RPKM) calculated from RNA sequencing results. All these four genes show significant differences between castes (false discovery rate < 0.05).

RNA-seq analysis showed that there were no chemoreceptor genes with significantly different expression levels between soldier and worker antennae. Notably, most chemoreceptor genes previously identified in *R. speratus* were incomplete, and full cDNA sequences were obtained for only 13–38% of these genes (ORs: 4/31, GRs: 9/25, IRs: 44/116). Full cDNA sequences were obtained for a larger percentage of non-receptor genes (60–67%; OBPs: 19/31, SNMPs: 3/5, CheAs: 2/3). Consequently, it is possible that the expression levels of chemoreceptor genes were not accurately obtained due to incomplete gene models. To definitively determine whether some chemoreceptor genes are differentially expressed between soldiers and workers, further endeavors should be undertaken to construct precise gene models based on long-read sequencing.

### Gene expression in soldier and worker antennae and other head parts

To verify the DEGs and confirm the specificity of gene expression in the antennae, qPCR analysis was performed. Soldiers and workers were obtained from each colony, and RNA samples were extracted separately from the antennae and other head parts (Fig. 3A). Based on the stability values obtained using GeNorm and NormFinder, we selected *EF-1alfa* as the internal control gene (Supplementary Table S10). Three non-receptor genes [*RsOBP27* (*RS009616*), *RsSNMP2* (*RS007977*), and *RsCheA1* (*RS004717*)] were significantly highly expressed in the antennae, especially in soldiers (Fig. 3B-D, a two-way analysis of variance (ANOVA) followed by Tukey’s test, *p* < 0.05). A two-way ANOVA was performed, and interactions were detected between castes (soldiers and workers) and body parts (antennae and other head parts) for all genes (*RsOBP27*: d.f. = 1, F = 13.6664, *p* = 0.00094; *RsSNMP2*: d.f. = 1, F = 16.8285, *p* = 0.00032; *RsCheA1*: d.f. = 1, F = 17.5847, p = 0.00025). Statistical differences between soldiers and workers were observed for all genes (*RsOBP27*: d.f. = 1, F = 12.917, *p* = 0.00123; *RsSNMP2*: d.f. = 1, F = 14.6724, *p* = 0.00066; *RsCheA1*: d.f. = 1, F = 19.4584, *p* = 0.00014). Significant differences were also found between body parts for all genes (*RsOBP27*: d.f. = 1, F = 65.2212, *p* = 8.6E-09; *RsSNMP2*: d.f. = 1, F = 57.2603, *p* = 3E-08; *RsCheA1*: d.f. = 1, F = 61.9569, *p* = 1.4E-08). We were not able to quantify the expression levels of *RsIR75o* (*RS012794*), because no PCR products were obtained using gene-specific primers (Supplementary Table S1). Thus, the RNA-seq results of three non-receptor genes [*RsOBP27* (*RS009616*), *RsSNMP2* (*RS007977*), and *RsCheA1* (*RS004717*)] were verified by independent qPCR analysis.

**Fig. 3.**
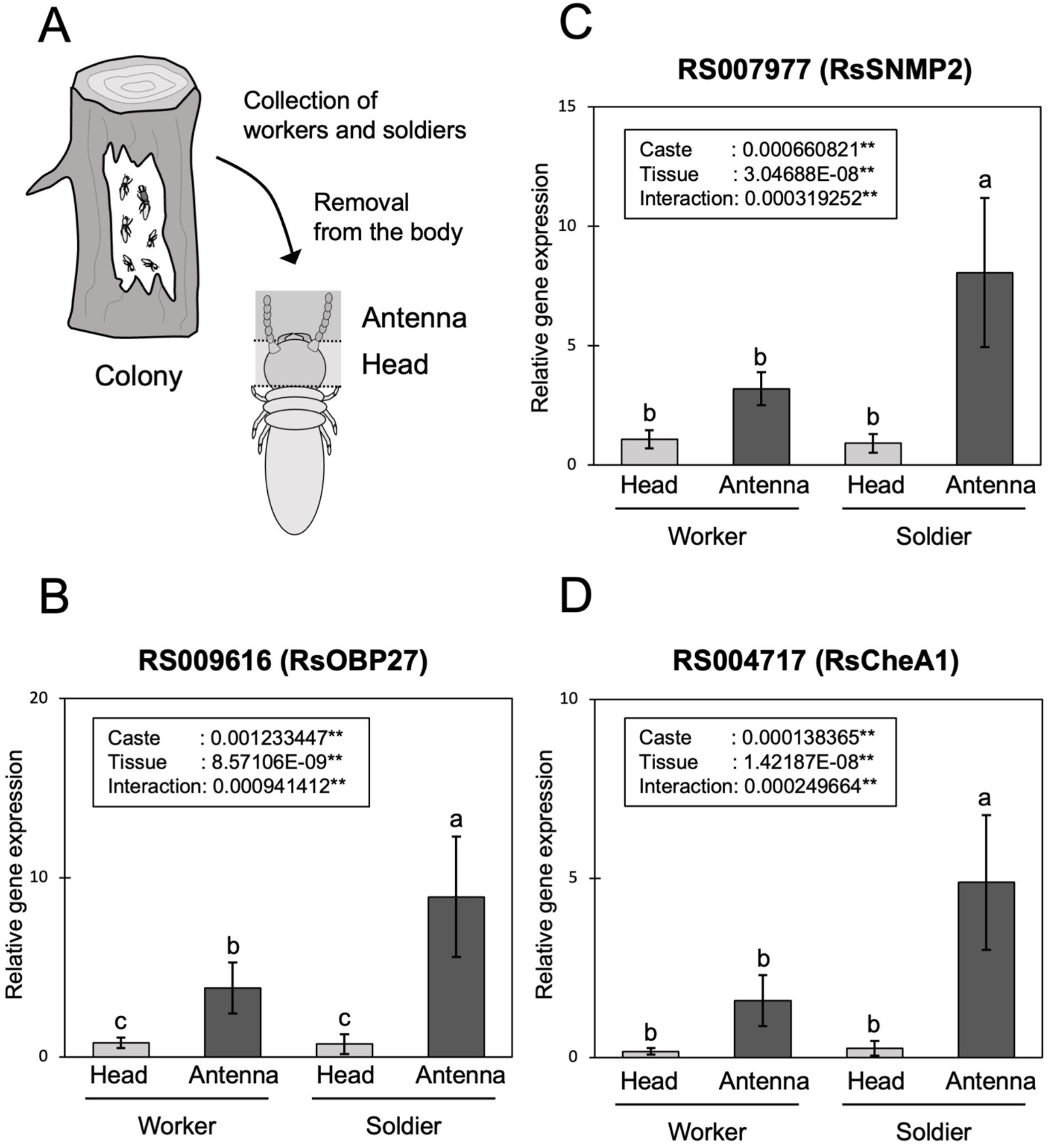
**(A)** Experimental design for sample preparation. **(B)**-**(D)** Expression levels of **(B)** *RS009616* (*RsOBP27*), **(C)** *RS007977* (*RsSNMP2*) and **(D)** *RS004717* (*RsCheA1*) in antennae and other head parts of *Reticulitermes speratus* soldiers and workers determined by qPCR. Each value was normalized to the expression levels of *EF-1alfa* (Supplementary Table S10). Different letters over the bars denote significant differences among the periods (two-way ANOVA followed by Tukey’s test, p < 0.05).

In some insect orders, including Lepidoptera, Hymenoptera, and Orthoptera, most *SNMP2* genes are specifically expressed in phalangeal cells of OSNs (Forstner et al., 2008; Zhang et al., 2015, Sun et al., 2019). These expression patterns suggest that SNMPs are involved in the reception of pheromones (Cassau & Kriegar, 2021). Moreover, antennal transcriptome analysis of the leafcutter ant, *Atta vollenweideri*, indicated that some *OBPs* have different expression patterns between castes and developmental stages (Koch et al., 2013). Gene expression analysis of antennal tissues of the fire ant, *Solenopsis invicta*, also showed that some *OBPs* are differentially expressed between castes (Zhang et al., 2016). In the parasitoid wasp, *Trichogramma japonicum*, gene expression levels were found to be significantly different between females and males (Wu et al., 2017). Although detailed gene function analyses have not been performed, especially for *Che* genes, these previous results suggest that several SNMPs and OBPs are involved in specific roles, including social and sexual behaviors.

The present results indicate that the expression levels of *RsOBP27* (*RS009616*), *RsSNMP2* (*RS007977*), and *RsCheA1* (*RS004717*) were higher in the soldier antennae than the worker antennae. These genes may be involved in soldier-characteristic (e.g., defensive and/or trophallactic) behaviors (discussed below). However, chemoreception genes with high expression levels in worker antennae were not identified in this study. The social tasks of termite workers are generally more complicated than those of soldiers (Roisin, 2000). In this study, 50 workers randomly chosen from the colonies were used to construct RNA-seq libraries. Consequently, it is possible that we failed to detect the expression patterns of chemoreception genes involved in worker-specific roles. For example, a previous study showed that two *CSP* genes (*RS000584* and *RS010442*) were significantly highly expressed in workers without soldiers compared to those with soldiers (using whole bodies without guts; Matsunami et al., 2022). However, we were unable to detect both of these genes in this study. To resolve these discrepancies, a comparison of gene expression patterns between workers engaged in different tasks (e.g., foraging, building, and nursing) is required.

### Gene expression in antennae of soldiers and workers isolated from colonies

To determine the different roles of the three DEGs identified, we performed qPCR analysis using individuals isolated from their own colonies at unusual caste ratios. Both soldiers and workers were isolated (15:15) for 3 days, and RNA samples were extracted from their antennae. Two-way ANOVA was performed, and interactions were detected between castes (soldiers and workers) and treatments (colony and isolation) for all genes (*RsOBP27*: d.f. = 1, F = 11.9896, *p* = 0.00174; *RsSNMP2*: d.f. = 1, F = 44.5641, *p* = 3E-07; *RsCheA1*: d.f. = 1, F = 20.102, *p* = 0.00011). Significant differences between soldiers and workers were observed for all genes (*RsOBP27*: d.f. = 1, F = 20.4003, *p* = 0.0001; *RsSNMP2*: d.f. = 1, F = 78.9041, *p* = 1.2E-09; *RsCheA1*: d.f. = 1, F = 77.2913, *p* = 1.52E-09). Significant differences were also found between treatments for all genes (*RsOBP27*: d.f. = 1, F = 9.38174, *p* = 0.0048; *RsSNMP2*: d.f. = 1, F = 106.489, *p* = 4.8E-11; *RsCheA1*: d.f. = 1, F = 9.80605, *p* = 0.00405). The expression levels were compared to those in soldiers and workers from the colonies (Fig. 4A). The results indicated that the expression patterns of soldier-biased chemoreception genes fluctuated significantly depending on the social context (Fig. 4B–D). Generally, soldier ratios tend to be relatively constant in each termite colony because both an excess and a lack of soldiers have negative effects on colony growth (Watanabe et al., 2014). The frequency of interactions between soldiers and workers may change in response to an increase or decrease in soldier ratios. We suggest that changes in the expression of chemoreception genes in soldier antennae are reflected in these behavioral changes among colony members.

**Fig. 4.**
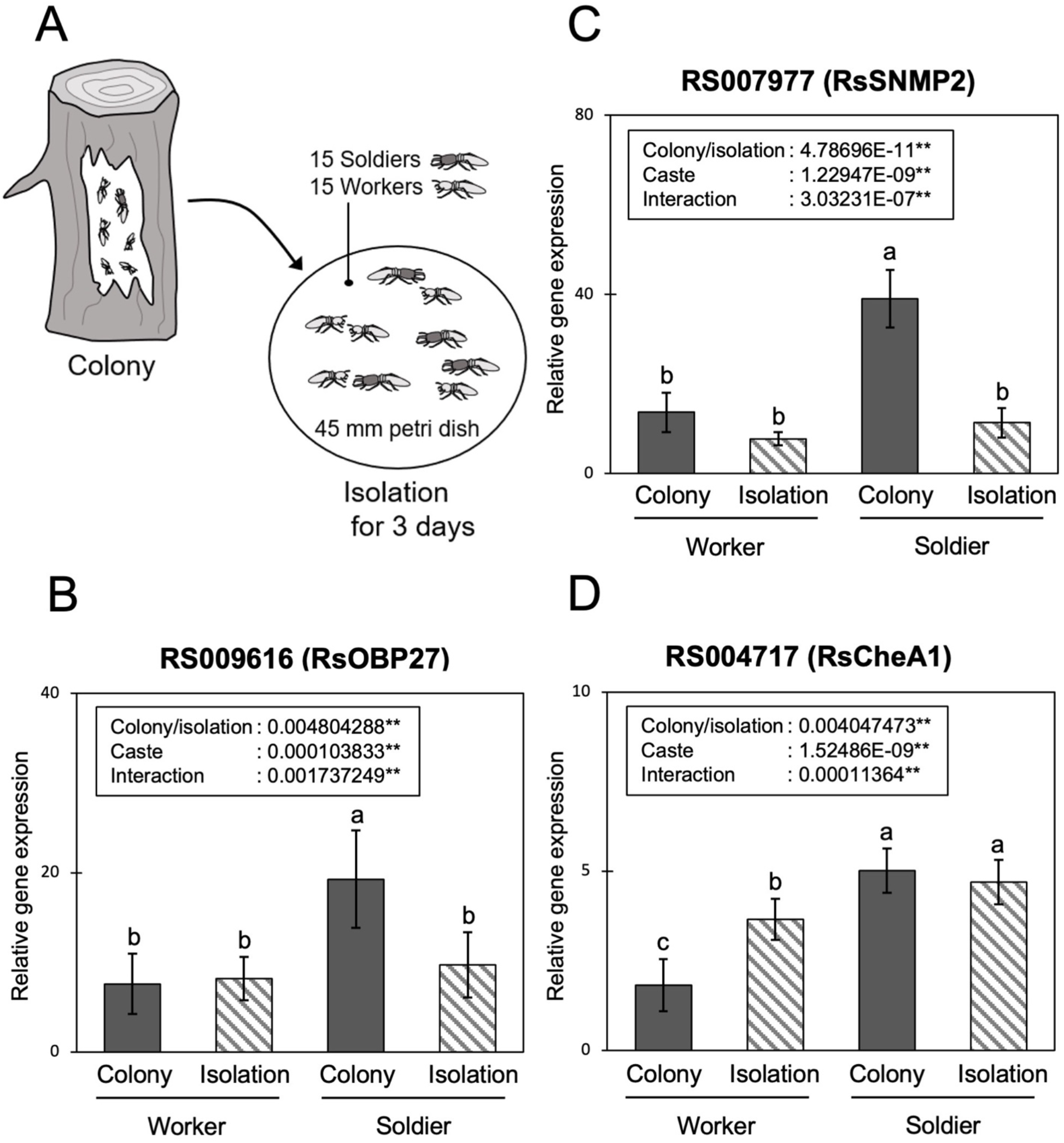
**(A)** Experimental design for sample preparation. **(B)**-**(D)** Expression levels of **(B)** *RS009616* (*RsOBP27*), **(C)** *RS007977* (*RsSNMP2*) and **(D)** *RS004717* (*RsCheA1*) in *Reticulitermes speratus* soldier and worker antennae determined by qPCR. Each value was normalized to the expression levels of *EF-1alfa* (Supplementary Table S10). Different letters over the bars denote significant differences among the periods (two-way ANOVA followed by Tukey’s test, p < 0.05).

Similar expression patterns were observed for *RsOBP27* (*RS009616*) and *RsSNMP2* (*RS007977*) (Fig. 4B–C). The highest expression levels were observed in the antennae of soldiers collected from their own colonies, and the expression levels significantly decreased after isolation from the colony. However, completely different expression patterns were observed for *RsCheA1* (*RS004717*; Fig. 4D). The expression levels in soldier antennae were consistently high, but those in worker antennae significantly fluctuated and increased after isolation from the colony. These different expression patterns suggest that *RsCheA1* (*RS004717*) has different roles or actions in soldier and worker antennae. Further detailed functional analyses are needed to confirm whether these three chemoreception genes are important for social behavior in *R. speratus*.

## Conclusion

We identified OBPs and CheAs in *R. speratus*, and performed RNA-seq analyses of worker and soldier (female and male) antennal tissues. The results showed that there were very few chemosensory genes with differential expression patterns between castes. We identified three non-receptor genes (*OBP, CheA*, and *SNMP*) with significantly high expression levels in soldier antennae. Independent qPCR analysis showed that the expression patterns of three genes in the soldier and worker antennae were changed in the different social contexts (different soldier ratios). These results suggest that several important chemoreception genes are likely involved in non-reproductive social behaviors in termites.

## EXPERIMENTAL PROCEDURES

### Characterization of RsOBPs and RsCheA

We first used blastp to search for gene models (Rspe OGS1.0; Shigenobu et al., 2022) of *R. speratus* for OBP and Che candidates using the protein sequences of other insect species (Vieira & Rozas, 2011; Terrapon et al., 2014; Robertson et al., 2018; Torres-Oliva et al., 2016) as queries with an e-value cutoff of 1.0E-5. We also ran a Hidden Markov Model (HMM) search using the PBP/GOBP family (pfam01395) and DUF1091 (pfam06477) as queries. We named the obtained candidates RsOBPs (31 genes) and RsCheAs (three genes).

### Phylogenetic analysis

We produced an alignment using the E-INS-i strategy of MAFFT (Kato et al., 2005) using the OBP protein sequences of *Drosophila melanogaster* [52 OBPs (Vieira & Rozas, 2011), 8 CheAs, and 13 CheBs (Torres-Oliva et al., 2016)], *Blattella germanica* [109 OBPs (Robertson et al., 2018) and 11 CheAs], *Zootermopsis nevadensis* [29 OBPs (Terrapon et al., 2014) and 5 CheAs], *R. speratus* (31 OBPs and 3 CheAs), and *Macrotermes natalensis* (21 OBPs and 4 CheAs). The OBPs of *M. natalensis*, and the CheAs of *B. germanica, Z. nevadensis*, and *M. natalensis* were newly identified using the same method used for *R. speratus*. Ambiguous sections of the alignment were removed using trimAl (gappyout option; Capella-Gutierrez et al., 2009). This alignment was used to produce a maximum likelihood phylogenetic tree using RAxML-NG (Kozlov et al., 2019) with 100 bootstrap replicates. The most appropriate models for amino acid sequence evolution [LG + I + G4 (OBP) and LG + I + G4 + F (Che)] were determined using ModelTest-NG (Darriba et al., 2019).

### Sample preparation and RNA-seq analysis

Three mature colonies were collected in 2017 from Furudo, Toyama Prefecture, Japan. Mature soldiers and W4-5 workers (6–7th instars; Takematsu, 1992; Maekawa et al., 2008) were collected from each colony. The sexes of individuals were identified based on the morphological characteristics of the 7–8th abdominal sternites (Zimet, 1982; Hayashi et al., 2003). Total RNA was extracted from the antennae (50 individuals) of soldiers and workers using an SV Total RNA extraction kit (Promega, Madison, WI, USA). The amount of RNA and DNA in each sample was quantified using a Qubit fluorometer (Life Technology, Eugene, OR, USA), and sample quality was confirmed using an Agilent 2100 bioanalyzer (Agilent Technologies, Palo Alto, CA, USA). Total RNA (500 ng) was used for cDNA synthesis and purification using a low-throughput protocol with a TruSeq Stranded RNA LT Kit (Illumina, San Diego, CA, USA). A half-scale reaction of the standard protocol was used for library preparation. The quality and quantity of cDNA were validated using an Agilent 2100 bioanalyzer and a KAPA qPCR SYBR Green PCR kit (GeneWorks, Thebarton, Australia). RNA-seq analysis was performed by single-end sequencing (66 bp) using a HiSeq 2500 instrument (Illumina, San Diego, CA, USA). Three replicates (i.e., biological triplicates) were prepared for both sexes of workers and soldiers, and 12 libraries were sequenced. All reads will be deposited in the DDBJ Sequence Read Archive database.

### Differentially expressed gene analysis

Prior to mapping the transcriptomic data, all libraries were processed to perform expression analysis as follows. First, the quality of the obtained sequence reads was determined using FastQC (Andrews, 2010) and the adaptor sequences were removed from all libraries using Cutadapt 1.4.2 (Martin, 2011) with the default parameters. Low-quality reads were trimmed using SolexaQA v2.5 (Cox et al., 2010) with a Phred score cutoff of 28 (-h 28) in DynamicTrim.pl and a minimum trimmed read length of 23 (-l 23) in LengthSort.pl. These reads were mapped to a *R. speratus* reference genome (gene model Rspe_OGS1.0; Shigenobu et al., 2022) using TopHat v2.0.9 (Kim et al., 2013) with the default parameters. Reads were counted using featureCounts v1.5.2 (Liao et al., 2014). The normalization factors for each library were calculated using the trimmed mean of M-values (TMM) method in the edgeR software package (Robinson and Oshlack, 2010). MA plots were used to represent the difference in the counts of each caste using edgeR. A false discovery rate < 0.05 was used as the cutoff for differential expression.

### Sample preparation for qPCR analysis

Three mature colonies were collected from Himi and Yatsuo, Toyama Prefecture, Japan, in 2021. The nest logs were brought back to the laboratory and kept in plastic containers at room temperature in constant darkness. Mature soldiers and W4-5 workers were collected from each colony (10 individuals of each caste) and used for RNA extraction, as described below. Dish assays were performed for each colony to determine the isolation effect. Filter paper (55 mm diameter; Advantec No. 1, Japan) was moistened with approximately 450 µL of distilled water and placed in a 65 mm Petri dish. A total of 15 workers were exposed to each filter paper with 15 soldiers, and all dishes were kept in an incubator at 25 °C in constant darkness for three days (one dish from each colony). All dishes were monitored daily and dead individuals were immediately removed from the dish. Ten workers and 10 soldiers were used for RNA extractions.

### qPCR analysis

Total RNA was extracted from the antennae of the soldiers and workers from each colony. The specimens were dissected on ice. The antennae were cut from 10 termites, and the other head parts were obtained from two termites (each sample from four colonies; four colonial replicates). Each tissue sample was immediately frozen in liquid nitrogen and then stored at -80 °C. Total RNA extraction and DNase treatment were performed using the ReliaPrep RNA Tissue Miniprep System (Promega). RNA and DNA were quantified using a Qubit 2.0 fluorometer. RNA purity and quantity were determined by spectroscopic measurements at 230, 260, and 280 nm using a NanoVue spectrophotometer (GE Healthcare Bio-Sciences, Tokyo, Japan). Single-stranded cDNA was synthesized from equal quantities of DNase-treated RNA (75 ng) using a High-Capacity cDNA Reverse Transcription Kit (Thermo Fisher, Waltham, MA, USA). We designed qRT-PCR primers for all three target genes using Primer3Plus (Untergasser et al., 2012; Supplementary Table S1). The relative quantification of transcripts was performed using the Thunderbird SYBR qPCR Mix (TOYOBO, Osaka, Japan) and a QuantStudio 3 Real-Time PCR System (Thermo Fisher). According to previous studies (Miyazaki et al., 2021; Yaguchi et al., 2022), the suitability of six reference genes, *beta-actin* (GenBank accession no. AB520714, Maekawa et al., 2010), *EF-1alfa* (AB602838, Hojo et al., 2011), *NADH-dh* (AB602837, Hojo et al., 2011), *eukaryotic initiation factor 1A* (EIF-1, gene ID: RS005199, Shigenobu et al., 2022), *glutathione-S-transferase 1* (GstD1, RS001168), and *ribosomal protein S18* (RPS18, RS015150) in *R. speratus* were evaluated using GeNorm (Vandesompele et al., 2002) and NormFinder (Andersen et al., 2004) software. A statistical test was performed using a two-way analysis of variance (ANOVA), followed by Tukey’s test, using the statistical software, Mac Statistical Analysis ver. 2.0 (Esumi, Tokyo, Japan).

## FIGURES

**Supplementary Fig. S1.**
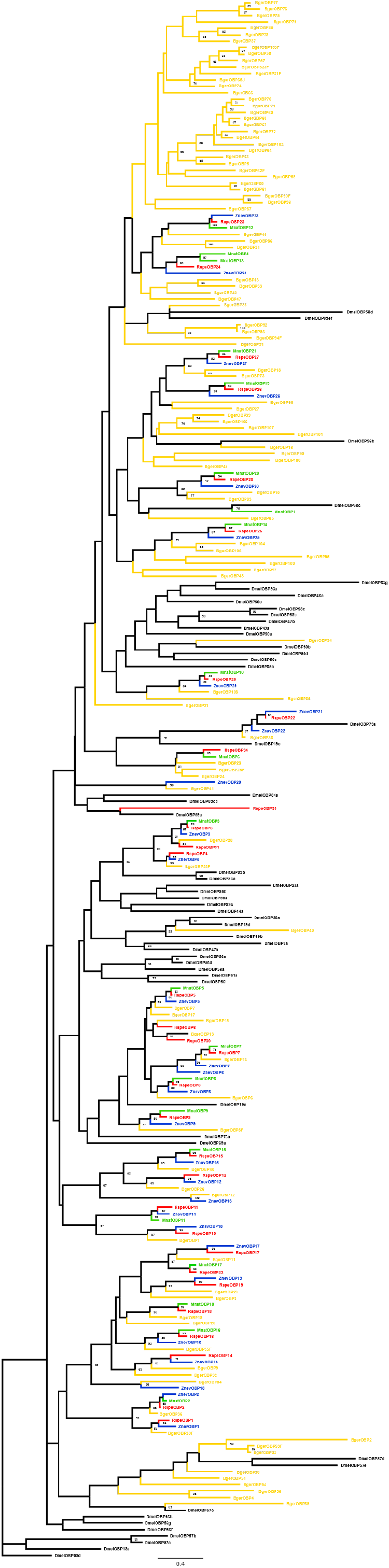
Maximum likelihood tree inferred from the amino acid sequences of OBPs identified from *Reticulitermes speratus* (Rsp), *Macrotermes natalensis* (Mnat), *Zootermopsis nevadensis* (Znev), *Blattella germanica* (Bger) and *Drosophila melanogaster* (Dmel). Bootstrap values (100 times) are shown for each node. Note that this is an unrooted tree.

**Supplementary Fig. S2.**
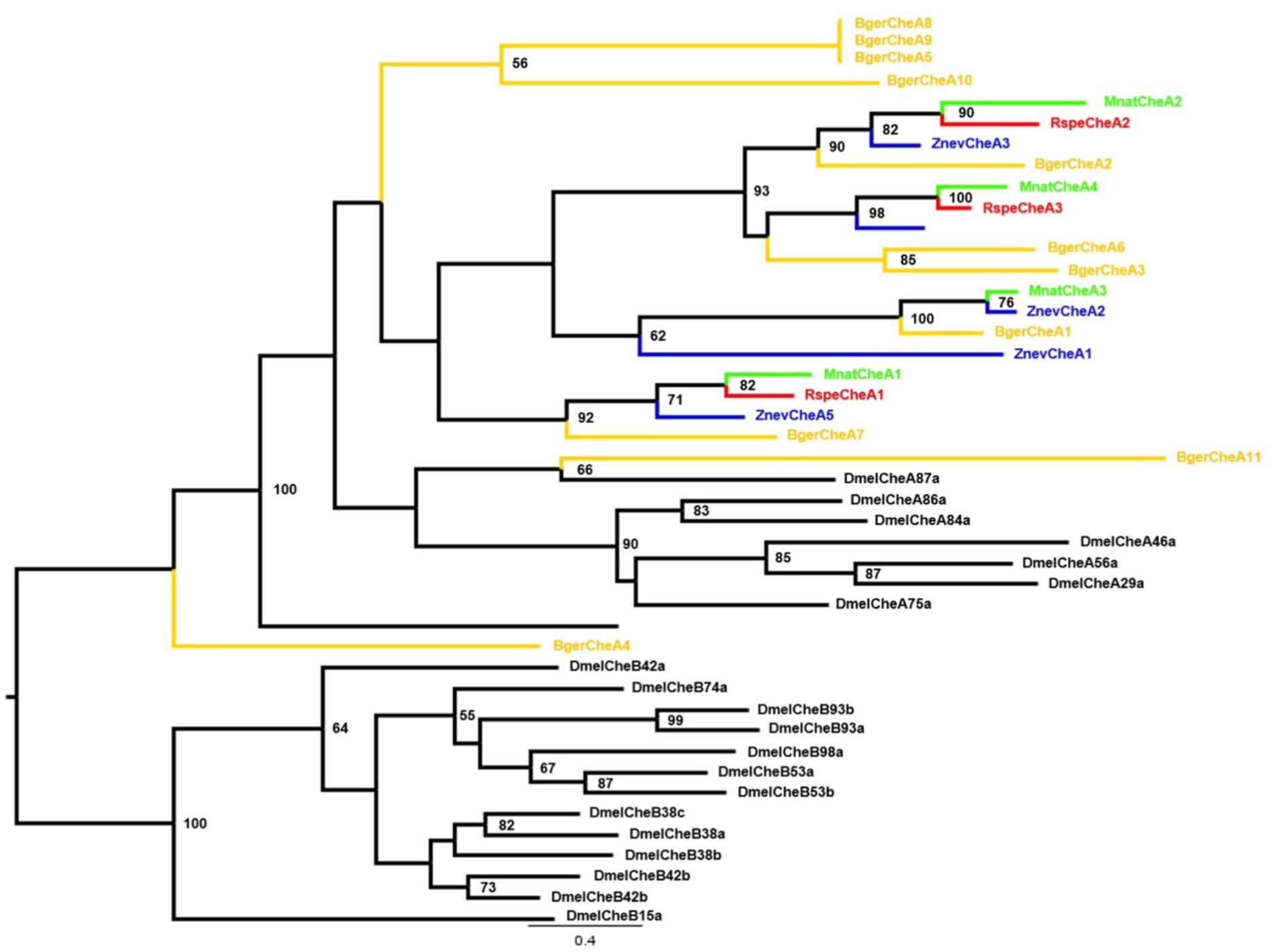
Maximum likelihood tree inferred from amino the acid sequences of Ches identified from *Reticulitermes speratus* (Rspe), *Macrotermes natalensis* (Mnat), *Zootermopsis nevadensis* (Znev), *Blattella germanica* (Bger) and *Drosophila melanogaster* (Dmel). Bootstrap values (100 times) are shown for each node. *Drosophila melanogaster CheB* (*DmelCheB*) genes were used as outgroups.

